# Structures, conformations and distributions of SARS-CoV-2 spike protein trimers on intact virions

**DOI:** 10.1101/2020.06.27.174979

**Authors:** Zunlong Ke, Joaquin Oton, Kun Qu, Mirko Cortese, Vojtech Zila, Lesley McKeane, Takanori Nakane, Jasenko Zivanov, Christopher J. Neufeldt, John M. Lu, Julia Peukes, Xiaoli Xiong, Hans-Georg Kräusslich, Sjors H.W. Scheres, Ralf Bartenschlager, John A.G. Briggs

## Abstract

Severe acute respiratory syndrome coronavirus 2 (SARS-CoV-2) virions are surrounded by a lipid bilayer from which spike (S) protein trimers protrude. Heavily glycosylated S trimers bind the ACE2 receptor and mediate entry of virions into target cells. S exhibits extensive conformational flexibility: it modulates the exposure of its receptor binding site and later undergoes complete structural rearrangement to drive fusion of viral and cellular membranes. The structures and conformations of soluble, overexpressed, purified S proteins have been studied in detail using cryo-electron microscopy. The structure and distribution of S on the virion surface, however, has not been characterised. Here we applied cryo-electron microscopy and tomography to image intact SARS-CoV-2 virions, determining the high-resolution structure, conformational flexibility and distributions of S trimers in situ on the virion surface. These results provide a basis for understanding the conformations of S present on the virion, and for studying their interactions with neutralizing antibodies.

## Introduction

Severe acute respiratory syndrome coronavirus 2 (SARS-CoV-2) is a betacoronavirus *(1,2)*, an enveloped virus containing a large nucleoprotein (N)-encapsidated positive sense RNA genome *(3)*. Three transmembrane proteins are incorporated into the viral lipid envelope: spike protein (S) and two smaller proteins, membrane protein (M) and envelope protein (E) *(3,4)*. When imaged by cryo-electron microscopy (cryo-EM), betacoronaviruses appear as approximately spherical particles, with variable diameters centered around 100 nm, containing a dense viroplasm, and bounded by a lipid bilayer from which prominent S trimers protrude *(5,6).* S trimers of SARS-CoV-2 identify and bind to the receptor, ACE2, on the surface of target cells and mediate subsequent viral uptake and fusion *(7–11)*. In doing so it undergoes a dramatic structural rearrangement from the prefusion form to the postfusion form *(12)*. The overall architectures of both pre and postfusion forms are well conserved among coronaviruses *(12–14)*.

During infection, S protein production takes place in an abnormal cellular environment. Coronaviruses extensively remodel the internal membrane organisation of the cell, generating viral replication organelles in which replication takes place *(15–17)*. The S protein, together with the other membrane protein M, and E, are inserted into membranes of the endoplasmic reticulum (ER), and traffic to the ER Golgi intermediate compartment (ERGIC). The encapsidated genome buds into the ERGIC to form virions which are then trafficked to the plasma membrane and released *(15–17)*.

The prefusion structure of S from coronaviruses including SARS-CoV-2 has been extensively studied using ectopic expression of a soluble, secreted form of S, followed by purification combined with cryo-EM and single particle reconstruction *(7,8,18–20)*. In the prefusion form, the receptor binding domain (RBD) sits at the top of a broad, trimeric spike, above the fusion core. Three copies of the RBD are surrounded by three copies of the N-terminal domain (NTD) which show some mobility *(7,8,12,18)*. In the closed prefusion conformation all three copies of the RBD lay flat on the spike surface, largely occluding the receptor binding site, while in the open prefusion conformation one or multiple RBDs can lift up to expose the receptor binding site *(7,8,18,19)*. The surface of the trimer is extensively glycosylated with 22 potential N-linked glycosylation sites on each monomer *(7,8,21)*. After receptor binding, structural transition of the prefusion to the postfusion form brings the fusion peptide and the transmembrane domain together at one end of a long, needle-like structure centered around a three-helix bundle *(12)*. Five N-linked glycans are regularly spaced along the length of the postfusion spike *(12)*.

Obtaining a full understanding of how S proteins function and how they interact with the immune system, requires knowledge of the structures, conformations and distributions of S trimers within virions. Here we have applied cryo-EM methods to study the structure, conformations and distributions of S trimers in situ on the virion surface.

## Results and discussion

In order to avoid artefacts associated with virus concentration or purification, we aimed to image SARS-CoV-2 virions from the supernatant of infected cells without virus concentration or purification. VeroE6 cells were infected with SARS-CoV-2 (isolate Germany/BavPat1/2020) *(22)*. At 48 h post-infection, supernatant was clarified, inactivated by fixation with formaldehyde and stored at −80°C. Western blot revealed that approximately 40 % of S protein on virions has been cleaved at the multibasic cleavage site into S1 and S2 (Fig. 1A). Fixed supernatant was vitrified by plunge freezing and imaged by cryo-EM. As expected, given the concentration of virus in cellular supernatants (around 10^7^ plaque forming units/ml), small numbers of individual virions were found scattered around the grid – these were imaged by cryo-electron tomography (cryo-ET) (Fig. 1B).

**Fig. 1.**
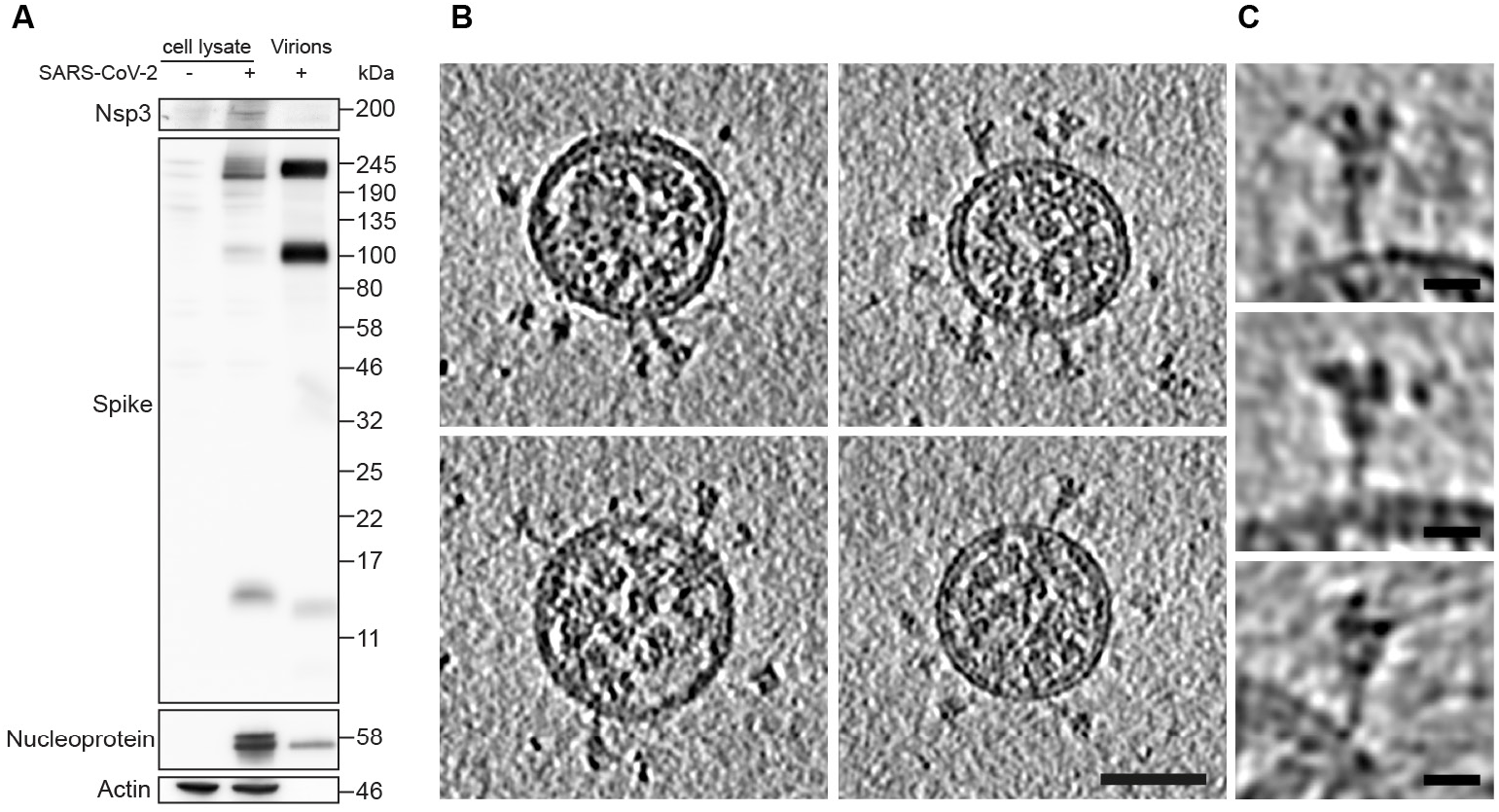
Characterization of virus production and representative images of intact, authentic SARS-CoV-2 virions. (**A**) Western blot analysis of SARS-CoV-2 S and N in lysates of VeroE6 cells and in virus preparations. In released virions, S is present in both cleaved and uncleaved forms. (**B**) Four representative tomographic slices of SARS-CoV-2 virions from the supernatant of infected cells. Virions are approximately spherical, contain granular density corresponding to N-packaged genome, and have S protein trimers protruding at variable angles from their surfaces. Scale bar 50 nm. (**C**) Three example S trimers from the dataset shown as projections through the trimer to illustrate the variable tilt towards the membrane. Scale bar 10 nm.

Virions were approximately spherical with a diameter to the outside of the lipid bilayer of 91 ± 11 nm (n=179) (Fig. S1A). They contain granular densities corresponding to the viral nucleoprotein, and are studded with S trimers (Fig. 1B,C). These features are generally consistent with those of other coronaviruses imaged by cryo-EM *(4–6)*. S trimers protruding from the viral surface had two morphologies – a minority were extended thin structures reminiscent of the postfusion form, while the majority are wider structures reminiscent of the prefusion form. This observation contrasts with a recent preprint showing cryo-EM images of purified SARS-CoV-2 virions inactivated with the nucleic acid modifier β-propiolactone in which only thin protrusions were seen on the viral surface *(23)*, and is consistent with *in situ* observations of virus assembly *(16)*.

We also collected tomograms of SARS-CoV-2 virions produced by infection of Calu-3 cells, a human lung carcinoma cell line that supports virus production to a titre comparable to Vero cells. The morphology of the virions and the appearance of the S trimers on the surface was consistent with that seen for virions produced from VeroE6 cells (Fig. S2).

Individual virions contained 25 ± 9 S trimers (Fig. S1B). A small sub-population of virions contained only few S trimers while larger virions contained more S trimers (Fig. S1B). We identified 4377 wide S trimers and 116 thin S trimers from 179 virions and subjected them to subtomogram averaging. The averaged structures, at 9 and 22 Å resolution, respectively, correspond very well to published structures of purified S trimers in the pre- and postfusion forms *(7,8,12)* (Fig. 2A). Overall, approximately 97 % of S trimers are in the prefusion form, and 3 % in the postfusion form. Pre- and postfusion forms appear to be distributed evenly among virions.

**Fig. 2.**
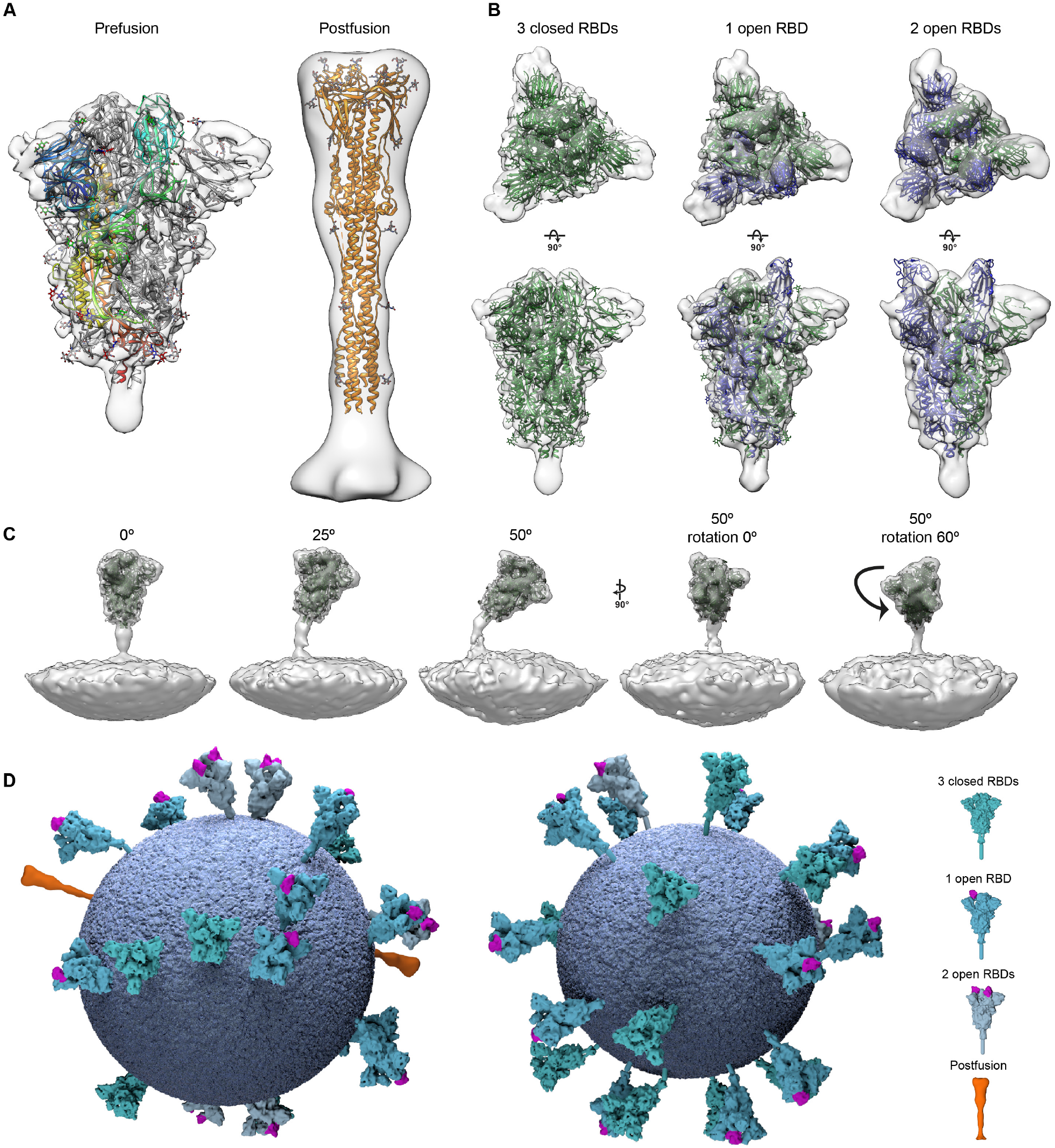
Structural analysis of SARS-CoV-2 S trimers from intact virions. (**A**) Structures of the prefusion (left) and postfusion (right) S trimer from intact virions determined by subtomogram averaging. The structures are shown as a transparent grey isosurface fitted with structures of the closed, prefusion SARS-CoV-2 S trimer (PDB 6VXX) and the postfusion SARS-CoV-1 S trimer (PDB 6M3W). In the prefusion form, one monomer is colored from blue (N terminus) to red (C terminus). The N-terminal domain is blue, the RBD appears cyan. Note that the NTD does not fully occupy the EM density because a number of loops are not resolved or built in PDB 6VXX. (**B**) The different conformations of the prefusion S trimer observed on intact virions by subtomogram averaging. Three conformations were observed: all RBDs in the closed position (left, fitted with PDB 6VXX); one RBD in the open position (center, fitted with PDB 6VYB); two RBDs in the open position (right, fitted with PDB 6X2B which does not include modelled glycans). Note that the two-open conformation has only been observed in vitro after inserting multiple stabilizing mutations. S monomers with closed RBDs are green, and with open RBDs are blue. (**C**) Averaging of subsets of trimers grouped according to their orientation relative to the membrane shows flexibility in the stalk region. Examples are shown for pools centered at 0°, 25° and 50° from the perpendicular, and for two different rotations of the trimer relative to the tilt direction. (**D**) 3D models of two individual SARS-CoV-2 virions with a membrane (blue) of the measured radius, and all spike proteins shown in the conformations, positions and orientations determined by subtomogram averaging. The different S conformations are distributed over the surface of the virions and can be tilted by up to ~60° relative to the membrane (Figure S1C).

To assess whether S trimers are present in open and/or closed conformations, we subjected the RBD regions of individual monomers within the trimers to classification. Three kinds of classes were found, those with the RBD in the closed position, those with the RBD in the open position, and those where the RBD was predominantly in the closed position, but with some weakening of the density, suggesting the presence of more mobile conformations (Fig. S3). Considering the classes to which each monomer was assigned, we derived structures of fully closed trimers, and of trimers where one RBD is open, both of which occur approximately equally frequently (~41% and ~45% of 4377 prefusion trimers) (Fig. 2B, Fig. S4). We also identified a small number of trimers (~13 % of 4377 prefusion trimers) in which two RBDs are in the open conformation (Fig. 2B). These observations confirm that the stochastic opening of the RBD observed in recombinant S trimers also takes place on the virus surface.

The trimers do not all protrude straight from the viral surface, but can tilt by up to 60° towards the membrane (Fig. S1C). We grouped trimers according to their orientation relative to the membrane, and averaged these groups independently. The averaged structures reveal that the membrane-proximal stalk region acts as a hinge with sufficient flexibility to allow tilting in all directions (Fig. 2C).

We generated models of individual virus particles, with S trimers located at the position, orientation and conformation that were determined by subtomogram averaging (Fig. 2D). S trimers appear to be distributed randomly on the viral surface, with no obvious clustering and no obvious relationship between location, orientation, and conformation. There is approximately one trimer per 500 nm^2^ of membrane surface compared to approximately one per 100 nm^2^ for influenza A *(24)*. The sparse distribution of S, together with the predominantly closed state, suggests that receptor binding may be less dependent on avidity effects than is the case for pandemic influenza viruses *(25)*.

The very low concentration of particles in cell supernatant makes it challenging to perform high-resolution structure determination. We therefore concentrated the virus by pelleting through a sucrose cushion. After concentration, the virus particles deviate from spherical morphology (Fig. S5), but overall features are preserved. We performed cryo-ET and subtomogram averaging on the particles, and saw predominantly prefusion S trimers, with occasional postfusion S trimers. Upon classification of the prefusion S we were only able to identify the RBD in the closed position, and monomers in which a weak RBD density is observed (Fig. S3C).

Virions in the supernatant of infected cells show primarily prefusion S trimers which are in either closed or open prefusion conformations. Virions concentrated through a sucrose cushion continue to show prefusion conformations, but the open conformation is no longer observed. Other studies have shown that virions fixed with β-propiolactone are primarily in a postfusion state *(23)*. We note that S trimers purified from membranes were found only in the closed prefusion and postfusion conformations *(12)*, while other studies have suggested that the open RBD in soluble S protein trimers is found in a continuum of different positions *(26)*. These observations suggest that the open prefusion conformations of the spike protein we observe in situ, but not after concentration, are fragile and may be affected by purification procedures, which has implications for vaccination strategies.

We next imaged the concentrated virus in 2D by cryo-EM and performed single-particle analysis on those prefusion S trimers that protruded from the sides of the virus particles as identified by a neural network picking algorithm, generating a consensus structure of the prefusion trimer at 3.7 Å resolution. Focussed classification with partial signal subtraction on individual RBD monomers led to two classes (Fig. S5). Consistent with the observations from cryo-ET, we observed 66% of the monomers in which the RBD is in the closed conformation and 34% of the monomers in which density for the RBD is weaker. We then refined the structures of S trimers in which all three RBDs are in the closed conformation, and those in which one RBD is weak, to resolutions of 3.9 Å and 4.3 Å, respectively (Fig. 3A, Fig. S6). The two structures are highly similar, differing only in the density levels for one RBD. We therefore used the structure with three closed RBDs to build and refine an atomic model. In this way we obtained an *in situ* structure for the S protein trimer on the viral surface.

**Fig. 3.**
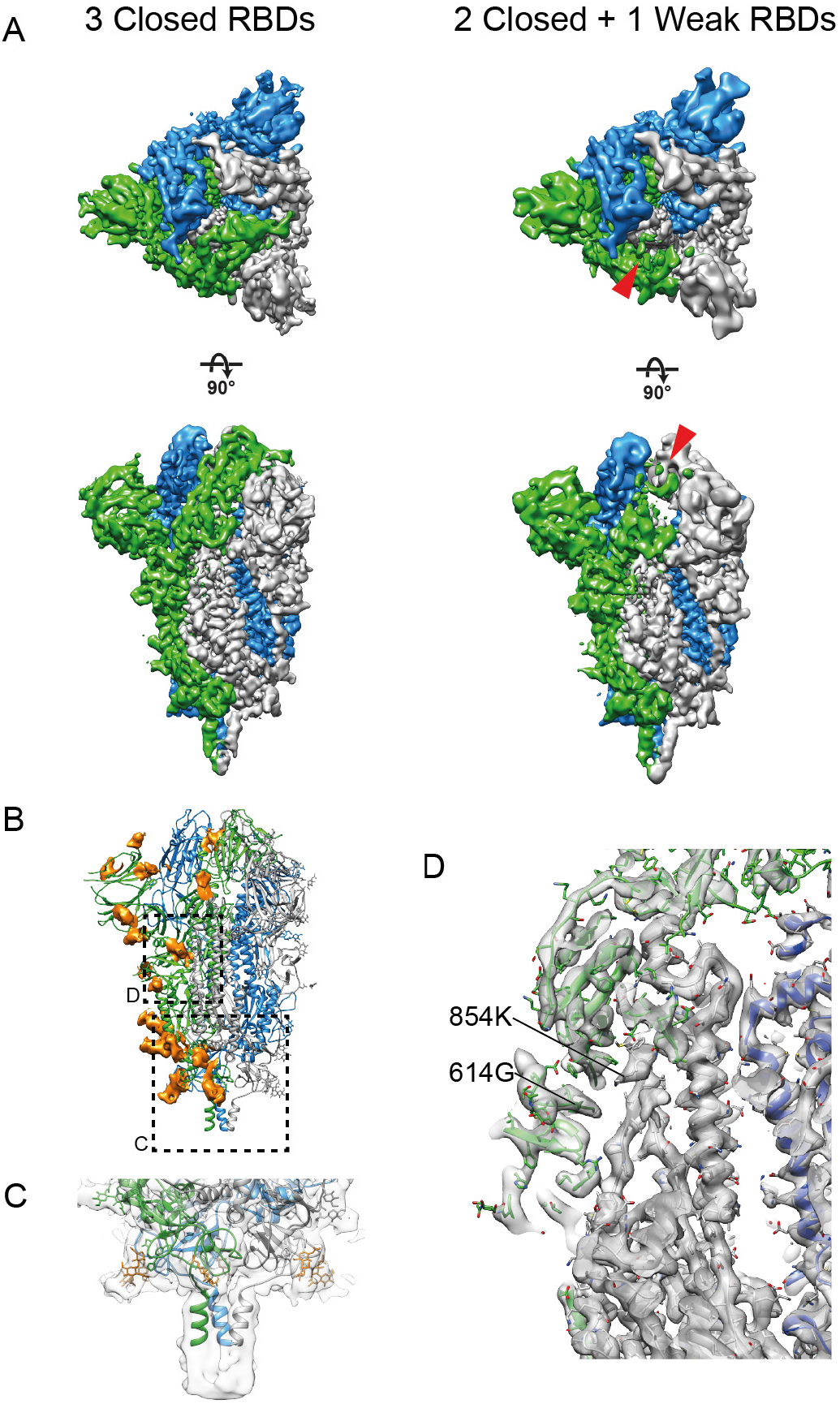
Structures of the SARS-CoV-2 S trimer determined on intact virions by single particle reconstruction. (**A**) Top and side views of S proteins with three closed RBDs (left, 3.9 Å resolution) and one weaker RBD (right, 4.3 Å resolution, weak RBD indicated by red arrowhead). Individual monomers are coloured white, blue and green. (**B**) Glycosylation profile of the S protein. Colour scheme as in (A), glycans are shown in orange. Boxes indicates the regions shown in C and D. (**C**) Close up of the base of the trimer at lower isosurface threshold to highlight the glycan ring and the extended C-terminal density. (**D**) Close up of the region of the spike where the D614G variation abolishes a salt bridge to 854K.

The positions of glycans on the surface of S are well resolved in our structure with strong density at 17 of the 22 predicted N-glycosylation sites (Fig. 3B). The other 5 glycosylation sites are in disordered NTD loops or in the stalk region not resolved at high resolution. In particular at the base of the trimer a clear ring of glycans forms a collar above the stalk region. The density for the stalk region extends by 2 helical turns compared to published structures of soluble ectodomain and further at lower resolution before fading out due to stalk flexibility (Fig. 3C). In contrast with the dense packing of haemagluttinin on influenza virions *(27)*, SARS-CoV-2 S trimers are sparsely distributed and can be highly tilted towards the membrane. Epitopes at the base of the head domain, and in the stalk region, would therefore be accessible to antibodies where they are not protected by the extensive glycan shell.

We compared the *in situ* structure of the S trimer to those previously obtained using exogenously expressed purified protein. A recent preprint described the structure of fulllength trimeric S solubilized in detergent micelles *(12)*. The study identified two features not present in most structures of soluble trimers of the S ectodomain: ordering of a loop between residues 833 and 853 (that is also ordered in the structure of the “locked” conformation of the ectodomain *(18);* and further ordering of the N-terminal residues to lock the RBDs in position. We observe weak density for N-terminal residues, and do not observe folded structure for the 833-853 region. The Germany/BavPat1/2020 SARS-CoV-2 strain we have imaged contains the widely circulating D614G substitution *(28)*, this mutation abolishes a salt bridge to K854 (Fig. 3D). We did not observe any additional density that would correspond to bound lipids such as those described in a recent preprint *(19)*, or other bound co-factors. These may be present sub-stoichiometrically or in rare conformations but are not a general feature of the S trimer *in situ.* Overall, our structure is very similar to that previously published for the soluble trimeric ectodomain in the closed prefusion form stabilized by a double proline mutation (Fig. S7) *(7,8)*. This provides an important validation of the use of recombinant, purified S protein trimers for research, diagnostics and vaccination. By determining the structure of S protein trimers to 3.7 Å resolution on the virion surface (Fig. 3), we demonstrate the potential of cryo-EM to allow the study of antibody binding to S in the context of the viral surface. This will be essential to understand how neutralizing antibodies, in particular against S2 and the stalk region, block virus infection, thus informing the design of prophylactic vaccines.

## Acknowledgements

We thank the staff of the MRC-LMB for generous support during the COVID-19 pandemic lockdown. We thank all the staff of the MRC-LMB EM Facility, in particular Anna Yeates, Grigory Sharov and Giuseppe Cannone, for supporting the EM experiments and Jake Grimmett and Toby Darling for supporting scientific computing. We are grateful to C. Drosten and EVAg for provision of the SARS-CoV-2 strain used here and we acknowledge microscopy support from the Infectious Diseases Imaging Platform (IDIP) at the Center for Integrative Infectious Disease Research Heidelberg. This study was supported by funding from the European Research Council (ERC) under the European Union’s Horizon 2020 research and innovation programme (ERC-CoG-648432 MEMBRANEFUSION to JAGB), the Medical Research Council as part of United Kingdom Research and Innovation (MC_UP_A025_1013 to SHWS; MC_UP_1201/16 to JAGB), and the Deutsche Forschungsgemeinschaft (240245660 – SFB 1129 to RB).

## Materials and methods

### Cells and Virus

VeroE6 cells were obtained from ATCC and were cultured in Dulbecco’s modified Eagle medium (DMEM, Life Technologies) containing 10% fetal bovine serum, 100 U/mL penicillin, 100 μg/mL streptomycin and 1% non-essential amino acids (complete medium). The Germany/BavPat1/2020 SARS-CoV-2 was isolated by Prof. Christian Drosten, Charité, Berlin, and distributed by the European Virology Archive (Ref-SKU: 026V-03883) at passage 2 (P2). A stock of SARS-CoV-2 was obtained by passaging the virus once in VeroE6 cells (P3). To produce SARS-CoV-2 virions, VeroE6 cells grown on 75 cm^2^ side-bottom tissue culture flasks were infected with SARS-CoV-2 (P3) at MOI of 0.5. Culture media from infected cells were harvested at 48 h postinfection, clarified by centrifugation at 1,000 × g for 10 min, cleared through a 0.45-μm nitrocellulose filter and fixed with 4% FA for 30 min at RT. Culture medium was supplemented with 10 mM HEPES (pH 7.2) prior to fixation. Virus-containing medium was subsequently aliquoted and stored at −80°C. Infectious supernatants containing SARS-CoV-2 virions were obtained from Calu-3 cells infected with P3 virus at an MOI = 5 for 48 h and processed as described above.

To obtain SARS-CoV-2 virions at high concentration, infection and harvest of VeroE6 culture medium were performed as above, followed by concentration of fixation-inactivated virions from media by ultracentrifugation through a 20% (wt/wt) sucrose cushion (120 min at 27,000 rpm in a Beckman SW32 rotor; Beckman Coulter Life Sciences). Pelleted particles were resuspended in PBS and stored in aliquots at −80°C.

### Western blot

VeroE6 or Calu-3 cells mock infected or infected for 48 h with SARS-CoV-2 (MOI = 5) were washed twice in PBS, scraped, pelleted at 700 x g for 5min and lysed in PBS pH 7.4, containing 1% Triton-X 100 and protease inhibitors (Merck) for 30 min at 4°C. Samples were centrifuged at 4°C for 30 min and supernatants were collected. Total protein concentration was calculated using the Bio-Rad Protein Assay kit (Biorad). Purified viruses were prepared for western blot by centrifugation of 32 ml of virus containing supernatants on a 10% sucrose cushion in a Beckmann J25 centrifuge. Centrifugation was performed at 10,000 x g for 4 h at 4°C. Supernatants were discarded and purified virus pellet were resuspended in 500 μl of PBS. For western blotting, 10 μg of total cell lysates and 5 μl of purified viruses were diluted in Laemmli buffer and loaded on a pre-casted Criterion XT 4-12% gradient gel (Biorad). Gels were transfered to PVDF membrane using a wet-electroblotting chamber system (Biorad) in Towbin buffer containing 10% methanol. Transfer was performed overnight at 4°C. Membranes were washed in PBS and blocked with 10% milk in PBS containing 0.2% tween-20 (PBS-T) for 1h. Membranes were incubated for 1 h at RT with primary antibody specific for spike (Abcam, cat# ab252690) or N-protein (Sino Biological, cat# 40143-MM05) both diluted at 1:1,000 in PBS-T. Next, the membranes were washed 3 times in PBS-T, incubated with HRP conjugated anti-mouse antibodies for 1 h, washed again 3 times in PBS-T, incubated with western Lightning Plus-ECL reagent (Perkin Elmer; Waltham, MA) and imaged using an Intas ChemoCam Imager 3.2 (Intas, Göttingen).

### RT-PCR and spike sequencing

Total RNA was isolated from infected VeroE6 cells 48 h after infection with Germany/BavPat1/2020 SARS-CoV-2 (passage 2). Spike cDNA was produced from the total RNA using superscript iii (ThermoFisher) with specific RT-primers (CAATTGTGAAGATTCTCATA). The cDNA was amplified by PCR using specific primers (Fwrd – ATG lllGlllllCTTG lllTATT; rev – TTATGTGTAATGTAAlllGA) and the resulting amplicon was sent for Sanger sequencing. Sequences were compared to the Germany/BatPat1/2020 SARS-CoV-2 reference sequence (Ref-SKU: 026V-03883) and found to be identical. Specific sequencing primers: Fwrd1 – ATGTTTGTTTTTCTTGTTTTATT; Fwrd2 – GGTTGGACAGCTGGTGCT; Fwrd3 – CCAACCATACAGAGTAGTAGTA; Rev1 – GTAGCAGCAAGATTAGCAGAA; Rev2 – TTATGTGTAATGTAATTTGA.

### Cryo-EM sample preparation

Virus samples from the supernatant of infected cells without any concentrations step (unconc) or concentrated by pelleting through a sucrose cushion (conc) were prepared, imaged, and processed in parallel. The virus suspension was mixed with 10-nm colloidal gold (in PBS solution) in 10:1 ratio. Then 3 μl of the solution was added to a glow-discharged copper grid (C-Flat 2/2, Protochips). Grids were plunge frozen into liquid ethane by backside blotting using a LeicaGP cryo plunger (Leica) and stored in liquid nitrogen until imaging.

### Cryo-electron tomography

Cryo-ET data collection was performed essentially as described previously *(29)*. Cryo-grids were loaded into an FEI Titan Krios transmission electron microscope operated at 300 kV and images were recorded on a Gatan K2 Summit direct detection camera in counting mode with a 20 eV energy slit in zero-loss mode. Tomographic tilt series between −60° and +60° were collected using SerialEM-3.8.0 software *(30)* in a dose-symmetric scheme *(31)* with a 3° angular increment. A total dose of 120 e^-^/A^2^ per tilt series was distributed evenly among 41 tilt images. The nominal magnification was 81,000 X, giving a pixel size of 1.532 Å on the specimen. The defocus range was between −2 μm and −6 μm and 10 frames were saved for each tilt angle. All data acquisition parameters are listed in Table S1.

Frames were motion-corrected in IMOD-4.10.30 *(32)* and images were dose-filtered using the *alignframes* function in IMOD 4.10.30. Exposure filtering was implemented according to the cumulative dose per tilt as described elsewhere *(33)*. The contrast transfer function (CTF) was measured using non-dose-filtered images using the ctfplotter package within IMOD *(34)*. Tilt series stacks were sorted using IMOD *newstack* function and fiducial-alignment of all tilt series was performed in IMOD/etomo. Tomograms with less than three trackable gold fiducials were discarded. Motion-corrected and dose-filtered tilt stacks were CTF-corrected by CTF multiplication and tomograms were reconstructed by weighted back-projected in novaCTF *(35)*. Tomograms were low-pass filtered to 50 Å for better visualization in EMAN2.2 *(36)* and tomographic slices were visualized with IMOD.

### Extraction of S trimers from tomograms

The initial steps of subtomogram alignment and averaging were implemented using MATLAB (MathWorks) scripts derived from the TOM *(37)* and AV3 *(38)* packages as described previously *(39,40)*. The missing wedge was modelled as the summed amplitude spectrum of background subtomograms for each tomogram, and was applied during alignment and averaging.

To generate an initial template model of the spike protein from the viral surface, 68 spikes were manually picked from four virions of tomograms that were down-scaled by 4x binning of the voxels. The 68 spikes’ initial Euler angles (2 out of 3) were determined based on the vector between two points, one on the head of the spike and one the membrane where the spike anchors, respectively. The 68 spikes were iteratively aligned to one another for four iterations applying three-fold symmetry to generate a low-resolution template that resembled a prefusion conformation of the spike. This template was used as an alignment reference for all virions (below). All postfusion spike proteins were manually identified and picked and initial Euler angles were assigned in the same manner.

The center of each virion was then marked manually using the Volume Tracer function in UCSF Chimera *(41)* and the radius of the virion was determined centered at the membrane using the Pick Particle Chimera Plugin *(42)*. An oversampled spherical grid of points was generated on the virion surface with ~9 nm spacing, and subtomograms were extracted for all grid points with a box size of 96 pixels (approximately 60 nm) centered at a radius 14 nm above these grid positions (approximately the radius of the expected center of the spike). Initial Euler angles were assigned to each subtomogram based on the orientation of the normal vectors relative to the sphere surface.

Subtomograms were aligned against the low resolution template using subTOM software. During this alignment, subtomogram positions converged onto clusters at the true spike positions. One subtomogram position was kept for each cluster, excluding particles within a distance of 10 pixels and removing particles with cross-correlation coefficients (CCC) below 0.09. Visual inspection of the tomograms using the Place Object Chimera Plugin *(42)* confirmed that the subtomograms selected in this manner corresponded to S trimers on the viral surface.

### Subtomogram averaging

Subsequent processing was performed in RELION *(43)*. For this purpose, subtomograms were reconstructed from the original tilt series images after motion correction using the relion_reconstruct program. Using dedicated python scripts, the S trimer positions in the 3D tomograms from the subTOM procedure outlined above were converted into 2D positions and defocus values in the corresponding tilt series images, as well as Euler angles in the RELION convention. Individual sub-tomograms were reconstructed at a 2x downscaled pixel size of 3.064 Å, by 3D insertion of Fourier slices of the cropped regions of the tilt series images, each multiplied by their corresponding CTF, which included the dose filter. Similarly, 3D-CTF volumes were generated by 3D insertion of the corresponding CTF^2^ slices. Subtomograms were reconstructed in a box size of 128 voxels for prefusion trimers and a box size of 192 voxels for postfusion trimers.

In order to deal with the CTF pre-multiplied sub-tomograms, as well as the multiplicity of each 3D voxel in the tilt series and the, a modified version of RELION was used for subtomogram refinement and classification (details to be described elsewhere). Standard 3D auto-refinement was performed with C3 symmetry and a soft-edged mask around the trimers, using a 30 Å low-pass filtered map generated from subTOM alignment parameters as initial reference. Using 4377 subtomograms, a 9 Å consensus map was calculated for the prefusion trimers; 116 postfusion trimers led to a 22 Å map.

Next, we performed symmetry expansion *(44)*, followed by focused classification without alignment and with partial signal subtraction, while keeping the orientations from the consensus refinement fixed. The mask used for focused classification was generated manually and enclosed the RBD of one monomer, including the closest NTD of the neighbouring monomer. Classification of the primary dataset (unconc1) led to three different RBD states: closed (50% of the monomers); open (25%) and with weak density (25%) (Fig. S3A). Classification of a second, smaller data set of virus particles in supernatant (unconc2), comprising 1566 trimers yielded 39% closed RBD, 24% open RBD and 38% weak RBD, in general agreement with the unconc1 results (Fig. S3B). Classification of a dataset of virus particles after concentration through a sucrose cushion (conc), comprising 3392 trimers, yielded 66% of the monomers corresponding to closed RBDs and 34% to RBDs with weak density. For this data set, no RBDs in the open conformation were identified (Fig S3C).

The classification of the RBDs as closed, open or with weaker density was used to divide the unconc1 dataset into three subsets: 41% (1789 trimers) have no open RBDs; 45% (1977 trimers) have one RBD in the open state; and 13% (572 trimers) have two open RBDs. The remaining 39 trimers have three RBDs in the open state and not further processed. In the class with weaker density, the RBD appears to be predominantly in the closed state and was treated as closed for this assignment. For each of the three subsets, reconstruction of the two independently refined half sets was performed using the orientations from the consensus refinement that gave the 9 Å consensus map described above. Subsequent standard post-processing procedures for resolution estimation, map sharpening and local-resolution filtering in RELION led to three final maps. The subset with no open RBDs gave a reconstruction with C3 symmetry in which all three RBDs were closed at 9.5 Å resolution. The other two subsets yielded structures with C1 symmetry and either one or two open RBDs, with resolutions of 9.3 Å and 10 Å, respectively (Figs. S3 and S4).

Averages of tilted spikes were generated by grouping according to the tilt and rotation of the subtomogram away from the normal vector to the membrane. Subtomograms were included in a group for averaging if they were within 15° of the displayed tilt (0°, 25° and 50°) and rotation (0°, 60°).

### Single particle Cryo-EM sample preparation and image acquisition

Virus solution concentrated through a 20% (wt/wt) sucrose cushion was frozen on C-Flat 2/2 3C grids (Protochips) following the same procedure as for cryo-ET, but without adding gold fiducials. Grids were imaged on a Thermo Fisher Scientific Titan Krios transmission electron microscope that was operated at 300 kV, using a Gatan K3 direct electron detector and a Gatan BioQuantum energy filter with a 20 eV energy slit. Movies with 48 frames and an accumulated dose of 50 electrons/Å^2^ were acquired in counting mode using SerialEM-3.8.0 *(30)* at a nominal magnification of 81,000 X, corresponding to a calibrated pixel size of 1.098 Å/pixel. Detailed data acquisition parameters are summarized in Table S2.

### Single particle Cryo-EM image processing

The Scheduler functionality in RELION-3.1 was used for fully automated real-time processing during data collection *(18,45)*. Movies were motion-corrected and dose-weighted using RELION’s implementation of the MotionCor2 algorithm *(46)*. Subsequently, dose-weighted sums were used to estimate the contrast transfer function (CTF) in CTFFIND-4.1.13 *(47)*. S trimers that were extending from the sides of virus particles were picked manually (973 particles from the first 100 micrographs) and then used as a training set for optimisation of the convolutional neural network in the automated particle picking software Topaz *(48)*. Extracted particles were subjected to 3D classification using a previously determined structure of the S trimer *(18)*, lowpass-filtered to 30 Å, as initial 3D reference. The selected 286,407 particles that contributed to 3D classes corresponding to S trimers were submitted to Bayesian polishing to correct for per-particle beam-induced motions and a second round of 3D classification to select the 55,159 particles that contributed to the best class. This final consensus set of particles was subjected to CTF refinement of per-particle defocus, per-micrograph astigmatism and beam tilt, followed by a second round of Bayesian polishing. 3D auto-refinements were performed with the selected particles after each round of 3D classification, CTF refinement or Bayesian polishing. The consensus structure had a resolution of 3.7 Å.

Subsequently, symmetry expansion *(44)*, followed by focused 3D classification with partial signal subtraction *(49)* was performed. Using a mask on a single RBD to focus classification into six classes, while keeping the orientations of the last consensus refinement fixed, resulted in the identification of two RBD states: closed or with weak density. S trimers with all three RBDs in the closed state were refined separately from S trimers with one RBD with weak density, resulting in two final maps with resolutions. Standard RELION post-processing was used for resolution estimation, map sharpening and local-resolution filtering. The C3 symmetric map with three closed RBD had an estimated overall resolution of 3.9 Å; the C1 map with one weaker RBD extended to 4.3 Å resolution.

### Single particle Cryo-EM model building and refinement

The SARS-CoV-2 S trimer structure (PDBID: 6VXX) was used as an initial model for building into the model with three closed RBDs (Fig S6, middle). Residues were adjusted manually in Coot 0.9 *(50)*. Steric clash and sidechain rotamer conformations were improved using the Namdinator web server. After further iterations of manual adjustment, the structure was refined in PHENIX-1.18.2 *(51)*. The geometry and statistics are given in Table S2. The unmasked model-to-map FSC was calculated in PHENIX for the refined model against the full reconstruction.

### 3D model building of spikes on authentic virions

In order to visualize the spike protein on the authentic SARS-CoV-2 virions, the coordinates, orientations and conformational classes determined by subtomogram averaging were converted into a format compatible with Maxon Cinema 4D, and imported together with the 3D models of the different conformational states determined by subtomogram averaging. The HR2 region was modeled as a cylinder. Images of individual virions from the dataset were rendered into Adobe Photoshop to generate images for presentation.

## Data availability

The cryo-EM and cryo-ET structures, and representative tomograms are deposited in the Electron Microscopy Data Bank (EMDB) under accession codes EMD-XXXXX – EMD-XXXXX. The associated molecular models are deposited in the protein data bank (PDB) under accession codes XXXX and XXXX.

---

**Fig. S1.**
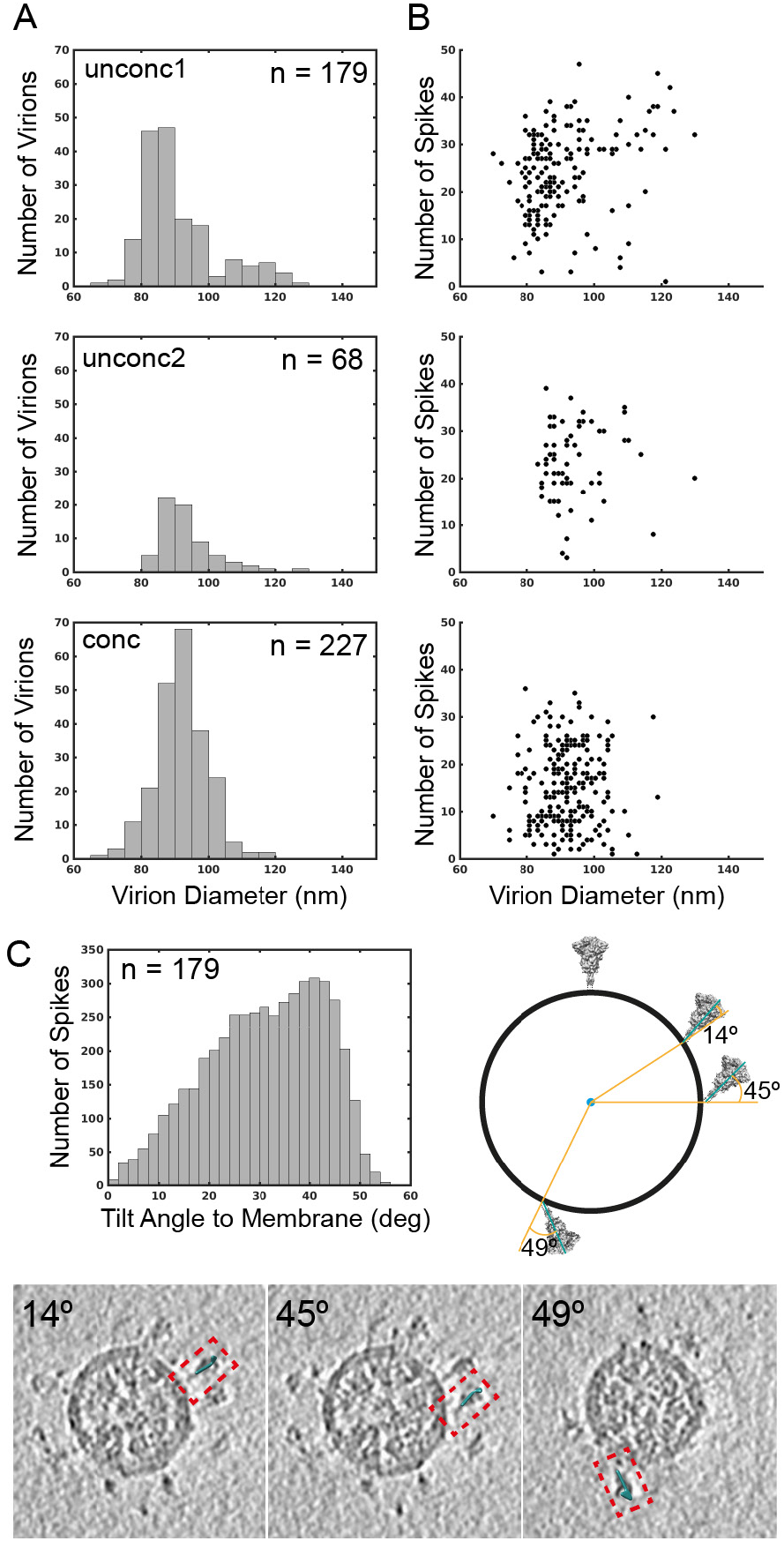
Characterization of SARS-CoV-2 virion morphology. (**A**) Histogram of virion diameters for unconcentrated extracellular virions in the supernatant of two independent preparations (top and middle), and for extracellular virions after concentration through a sucrose cushion (bottom). After concentration the virions become less spherical. Mean and standard deviation for diameters are 91 ± 11 nm (n=179), 94 ± 9 nm (n=68) and 92 ± 8 nm (n=227) for the three preparations. (**B**) Scatter plot of number of spikes identified per virion during subtomogram averaging against virion diameter for the same virions shown in panel (**A**). Visual inspection indicates that almost all spikes were identified for virions in the supernatant, but that not all spikes are identified in the concentrated preparation leading to an underestimate of the number of spikes. (**C**) Histogram of spike tilt angle towards the membrane for the larger supernatant virus dataset (unconc1). The schematic indicates the angle that was measured. Three examples of individual tilted spikes are marked on tomographic slices through an intact virion, with their associated angle.

**Fig. S2.**
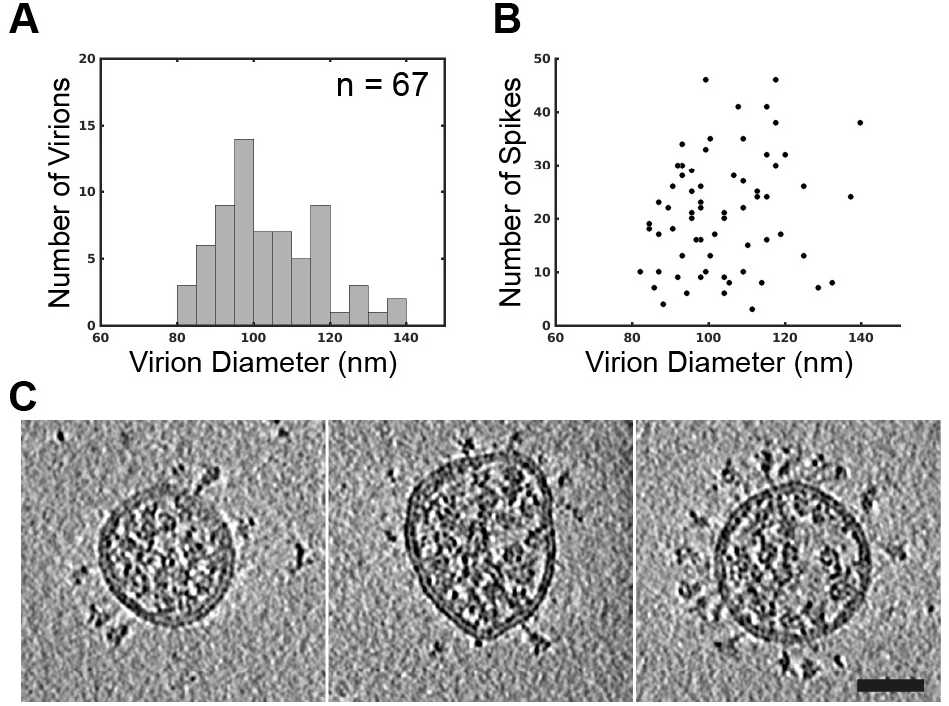
Morphology of SARS-CoV-2 virions released from infected Calu-3 cells. As in figure S1, (**A**) Histogram of virion diameters. Mean and standard deviation for diameters are 104 ± 13 nm (n=67) (**B**) Scatter plot of number of spikes identified per virion during subtomogram averaging against virion diameter for the same virions shown in panel A. (**C**) central slices through representative viruses. Virions from Calu-3 cells had a slightly broader diameter distribution than those from VeroE6 cells. Scale bar 50 nm. (**D**) Western blot analysis of SARS-CoV-2 S and N in cell lysates and in virus preparations. In released virions, S is present in both cleaved and uncleaved forms.

**Fig. S3.**
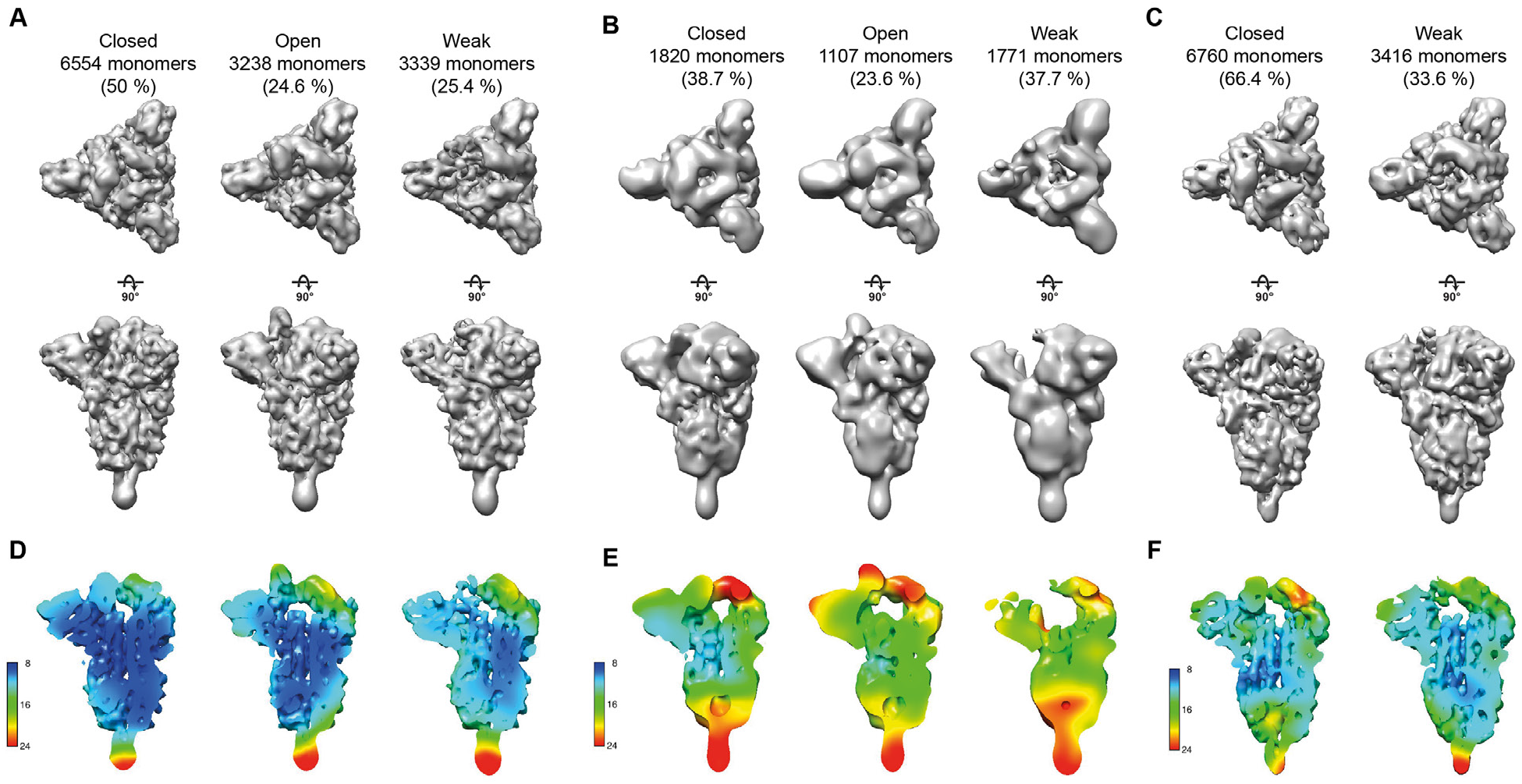
Classification of SARS-CoV-2 spike RBDs. (**A**) Class averages obtained after focussed classification on the RBD of the left monomer after symmetry expansion of the unconc1 dataset. Top views and side views are shown for closed, open and weak classes. (**B**) Equivalent analysis for a smaller, independent dataset (unconc2). (**C**) Equivalent analysis for a dataset obtained after concentrating virus through a sucrose cushion (conc). Only closed and weak classes were obtained. (**D-F**) Local resolution maps for structures shown in (A-C).

**Fig. S4.**
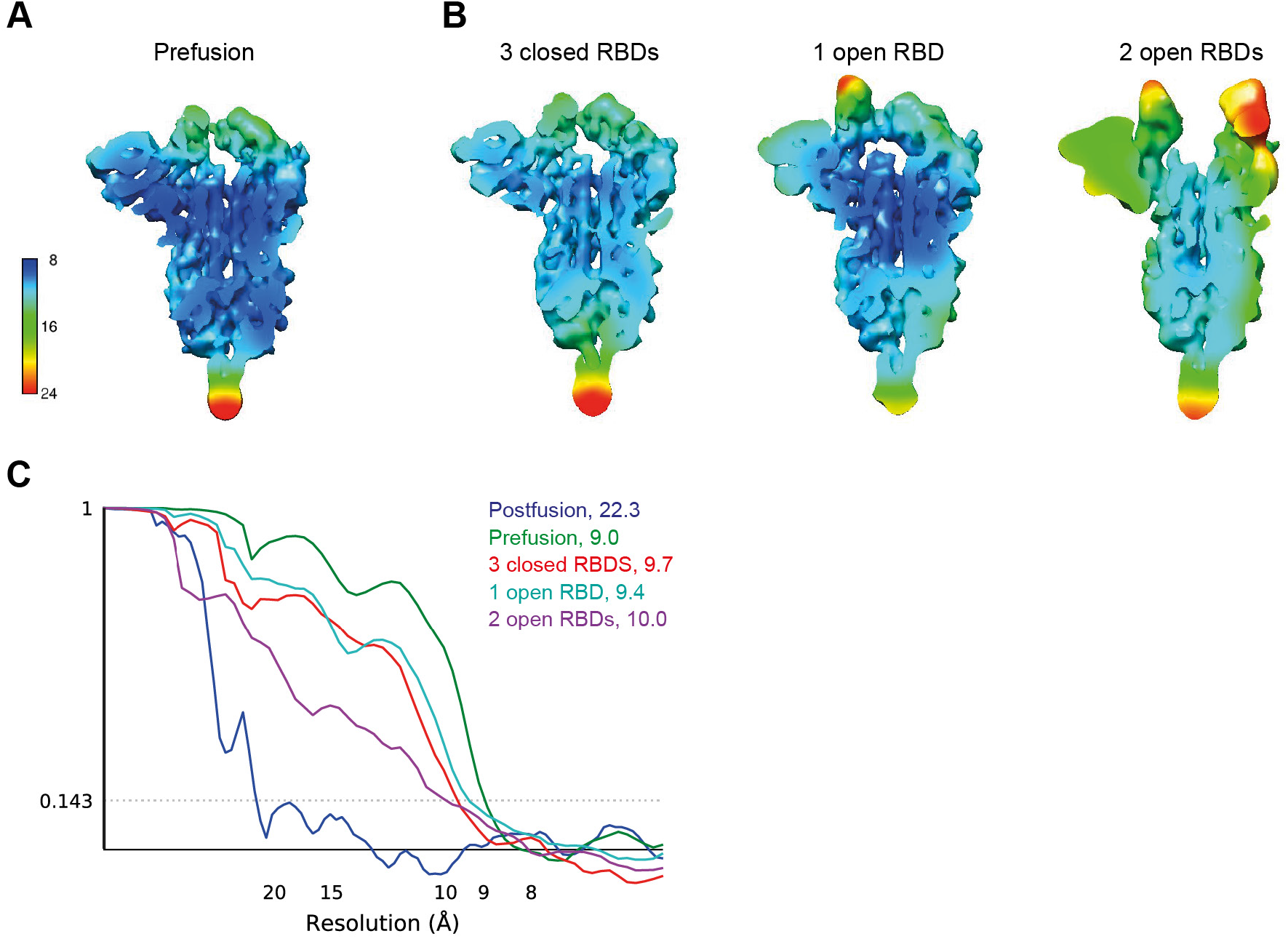
Resolution assessment of subtomogram averaging structures. (**A**) Local resolution map for the consensus structure obtained for the prefusion S trimers. (**B**) Local resolution maps for the prefusion S trimer in three different conformations. (**C**) Global resolution assessment by Fourier shell correlation at the 0.143 criterion for the four structures shown in A and B, as well as the postfusion S trimer.

**Fig. S5.**
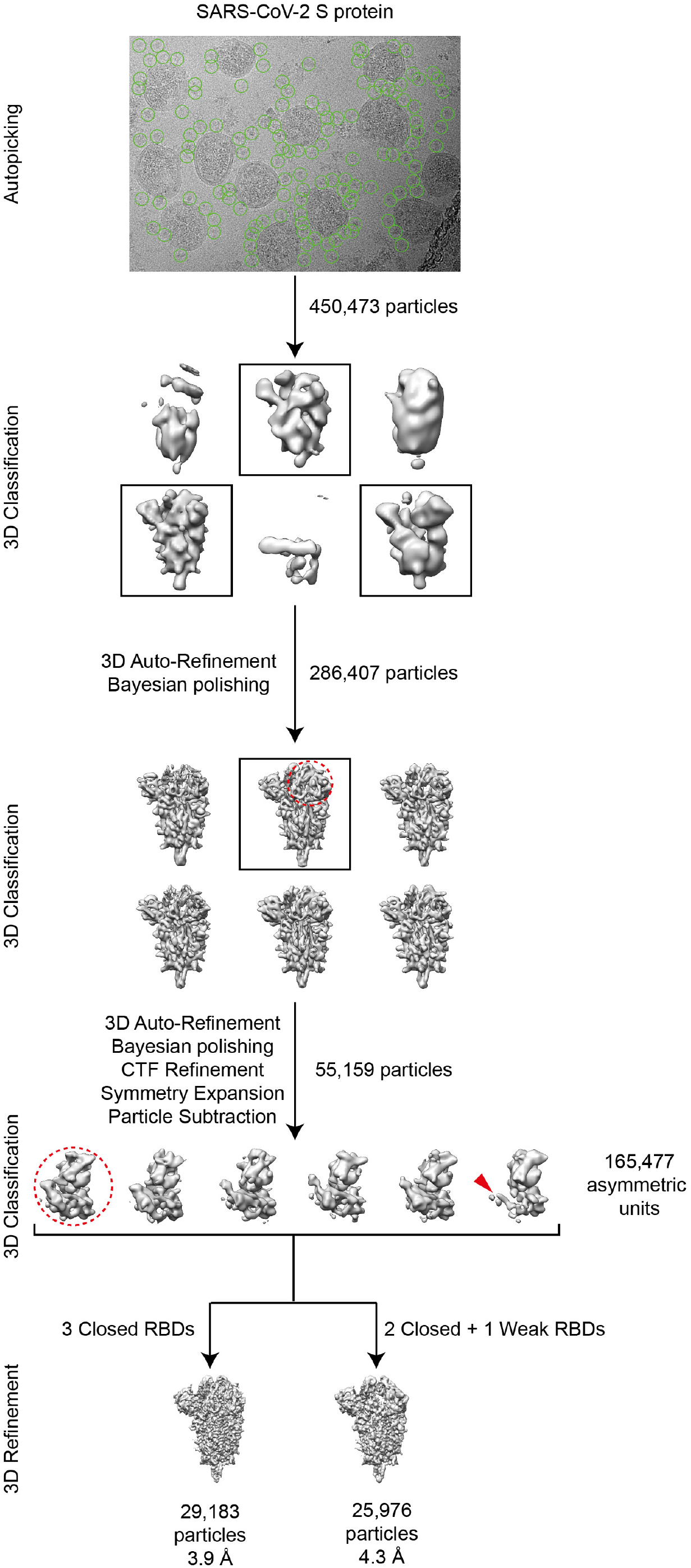
Single particle cryo-EM image processing workflow. Automatically picked particles (green circles) were subjected to 3D classification. Selected 3D classes are indicated by black boxes. RBDs from individual asymmetric units from the S trimer (red dashed circle) were locally classified to sort different conformations of RBD. S trimers with all three RBDs in the closed state were further refined with C3 symmetry. S trimers where one RBD had weak density (red arrowhead) were refined with C1 symmetry. For further details see materials and methods.

**Fig. S6.**
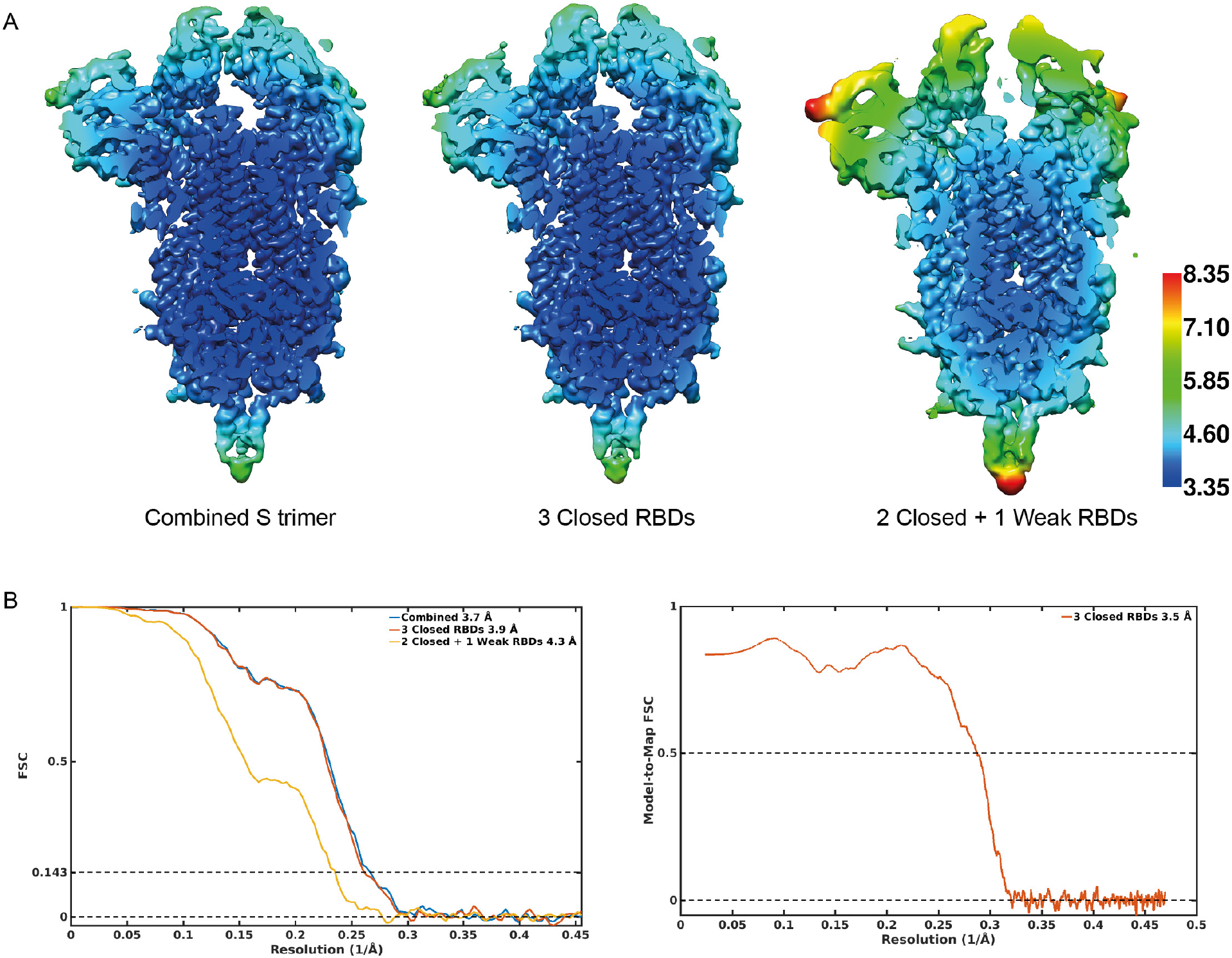
Single particle Cryo-EM structure validation. (**A**) Cut-open cryo-EM maps obtained using all prefusion S trimers, S trimers with 3 closed RBDs or S trimers with 2 closed and 1 weak RBDs, coloured according to the local resolution. (**B**) FSC curves for the three structures in (A), and for the atomic model against the map.

**Fig. S7.**
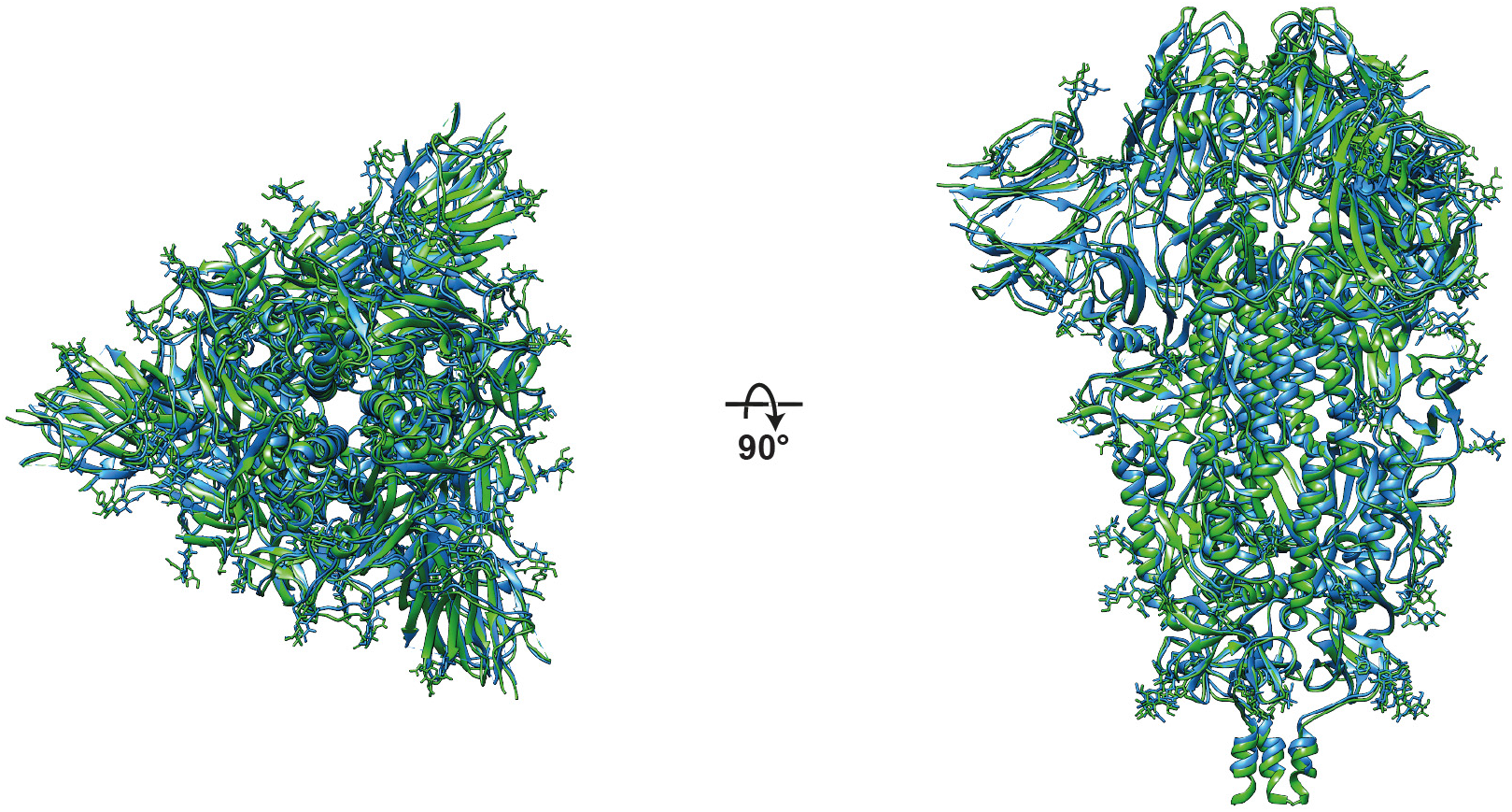
Structural comparison of in situ structure with recombinant soluble structure. Structural superposition of S trimer modelled into the structure of the trimer with three closed RBDs (green, this study) with the published structure of recombinant, soluble closed trimer (blue, PDB 6VXX). Top and side views are shown. The structures are very similar.

**Table S1.**
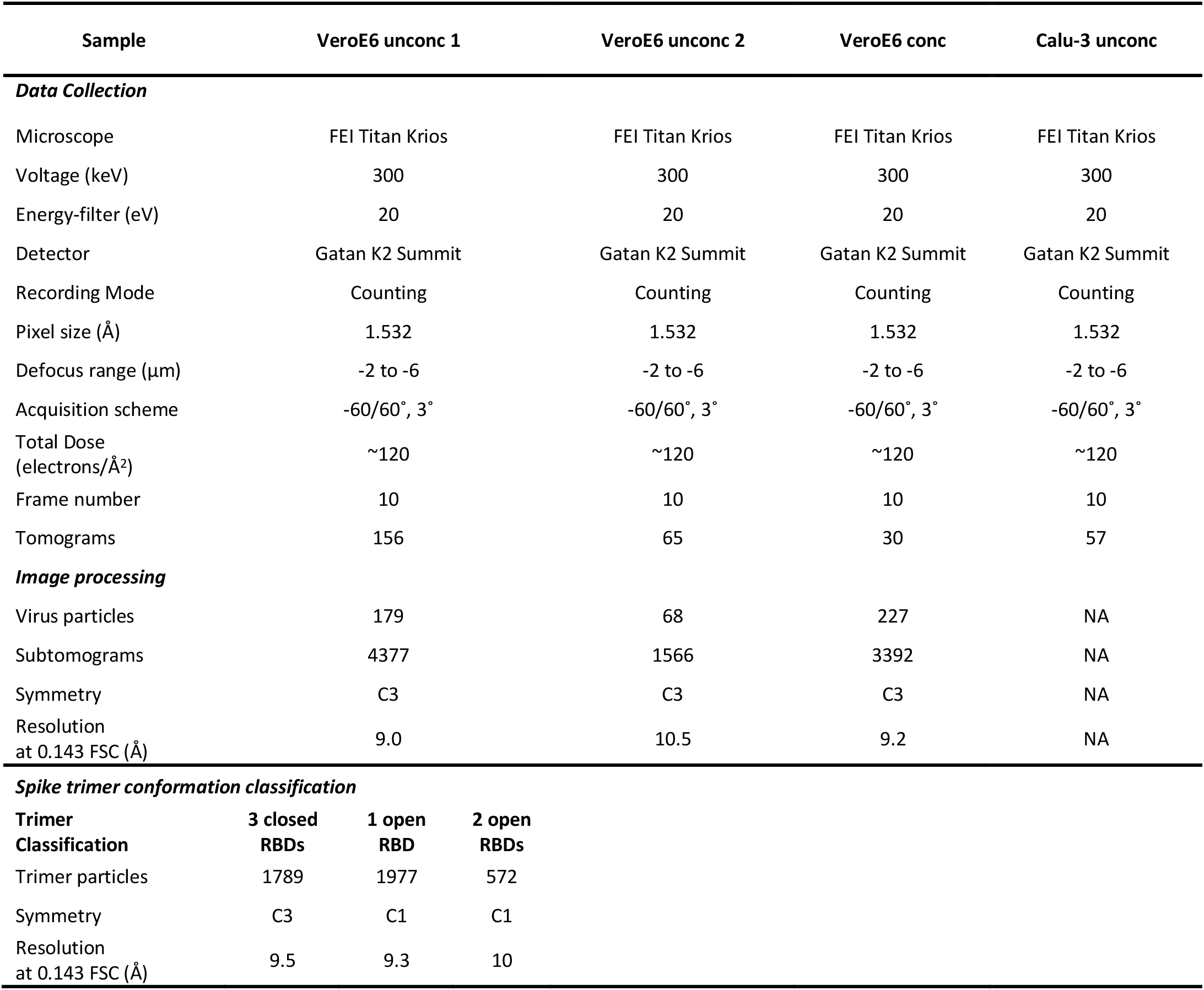
Cryo-ET data acquisition and image processing.

| Sample | VeroE6 unconc 1 | VeroE6 unconc 2 | VeroE6 conc | Calu-3 unconc |
| --- | --- | --- | --- | --- |
| **Data Collection** |  |  |  |  |
| Microscope | FEI Titan Krios | FEI Titan Krios | FEI Titan Krios | FEI Titan Krios |
| Voltage (keV) | 300 | 300 | 300 | 300 |
| Energy-filter (eV) | 20 | 20 | 20 | 20 |
| Detector | Gatan K2 Summit | Gatan K2 Summit | Gatan K2 Summit | Gatan K2 Summit |
| Recording Mode | Counting | Counting | Counting | Counting |
| Pixel size (Å) | 1.532 | 1.532 | 1.532 | 1.532 |
| Defocus range (µm) | -2 to -6 | -2 to -6 | -2 to -6 | -2 to -6 |
| Acquisition scheme | -60/60°, 3° | -60/60°, 3° | -60/60°, 3° | -60/60°, 3° |
| Total Dose (electrons/Å²) | ~120 | ~120 | ~120 | ~120 |
| Frame number | 10 | 10 | 10 | 10 |
| Tomograms | 156 | 65 | 30 | 57 |
| **Image processing** |  |  |  |  |
| Virus particles | 179 | 68 | 227 | NA |
| Subtomograms | 4377 | 1566 | 3392 | NA |
| Symmetry | C3 | C3 | C3 | NA |
| Resolution at 0.143 FSC (Å) | 9.0 | 10.5 | 9.2 | NA |
| **Spike trimer conformation classification** |  |  |  |  |
| **Trimer Classification** | **3 closed RBDs** | **1 open RBD** | **2 open RBDs** |  |
| Trimer particles | 1789 | 1977 | 572 |  |
| Symmetry | C3 | C1 | C1 |  |
| Resolution at 0.143 FSC (Å) | 9.5 | 9.3 | 10 |  |

**Table S2.**
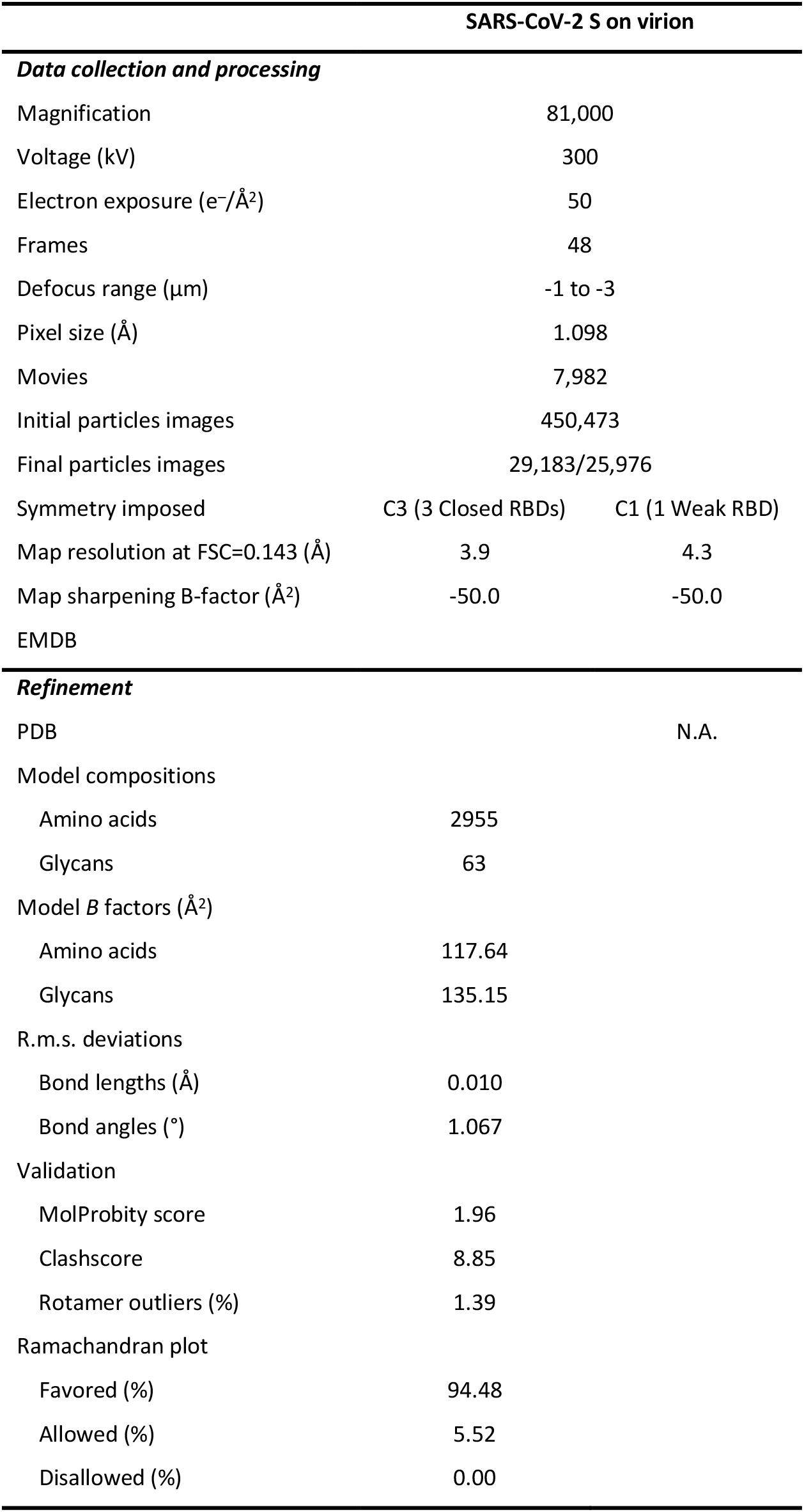
Cryo-EM data collection and refinement statistics

## References

1. Zhu, N. et al. A novel coronavirus from patients with pneumonia in China, 2019. N. Engl. J. Med. 382, 727–733 (2020).

2. Zhou, P. et al. A pneumonia outbreak associated with a new coronavirus of probable bat origin. Nature 579, 270–273 (2020).

3. Masters, P. s. & Perlman, S. CHAPTER 28 – Coronaviridae. in Fields Virology, 6th Edition vol. 1 826–858 (Elsevier, 2013).

4. Neuman, B. W. & Buchmeier, M. J. Supramolecular Architecture of the Coronavirus Particle. in Advances in Virus Research vol. 96 1–27 (2016).

5. Neuman, B. W. et al. Supramolecular Architecture of Severe Acute Respiratory Syndrome Coronavirus Revealed by Electron Cryomicroscopy. J. Virol. 80, 7918–7928 (2006).

6. Bárcena, M. et al. Cryo-electron tomography of mouse hepatitis virus: Insights into the structure of the coronavirion. Proc. Natl. Acad. Sci. U. S. A. 106, 582–587 (2009).

7. Wrapp, D. et al. Cryo-EM structure of the 2019-nCoV spike in the prefusion conformation. Science (2020) doi:10.1126/science.abb2507.

8. Walls, A. C. et al. Structure, Function, and Antigenicity of the SARS-CoV-2 Spike Glycoprotein. Cell 181, 281–292.e6 (2020).

9. Hoffmann, M. et al. SARS-CoV-2 Cell Entry Depends on ACE2 and TMPRSS2 and Is Blocked by a Clinically Proven Protease Inhibitor. Cell 181, 271–280.e8 (2020).

10. Shang, J. et al. Structural basis of receptor recognition by SARS-CoV-2. Nature 581, 221–224 (2020).

11. Wang, Q. et al. Structural and Functional Basis of SARS-CoV-2 Entry by Using Human ACE2. Cell 181, 894–904.e9 (2020).

12. Cai, Y. et al. Distinct conformational states of SARS-CoV-2 spike protein. bioRxiv 2020.05.16.099317 (2020) doi:10.1101/2020.05.16.099317.

13. Walls, A. C. et al. Tectonic conformational changes of a coronavirus spike glycoprotein promote membrane fusion. Proc. Natl. Acad. Sci. U. S. A. 114, 11157–11162 (2017).

14. Yuan, Y. et al. Cryo-EM structures of MERS-CoV and SARS-CoV spike glycoproteins reveal the dynamic receptor binding domains. Nat. Commun. 8, 15092 (2017).

15. Fehr, A. R. & Perlman, S. Coronaviruses: An overview of their replication and pathogenesis. Methods Mol. Biol. 1282, 1–23 (2015).

16. Klein, S. et al. SARS-CoV-2 structure and replication characterized by in situ cryoelectron tomography. bioRxiv 2020.06.23.167064 (2020) doi:10.1101/2020.06.23.167064.

17. Snijder, E. J. et al. A unifying structural and functional model of the coronavirus replication organelle: Tracking down RNA synthesis. PLOS Biol. (2020) doi:10.1371/journal.pbio.3000715.

18. Xiong, X. et al. A thermostable, closed, SARS-CoV-2 spike protein trimer. bioRxiv 2020.06.15.152835 (2020) doi:10.1101/2020.06.15.152835.

19. Toelzer, C. et al. Unexpected free fatty acid binding pocket in the cryo-EM structure of SARS-CoV-2 spike protein. bioRxiv 2020.06.18.158584 (2020) doi:10.1101/2020.06.18.158584.

20. Henderson, R. et al. Controlling the SARS-CoV-2 Spike Glycoprotein Conformation. bioRxiv 2020.05.18.102087 (2020) doi:10.1101/2020.05.18.102087.

21. Watanabe, Y., Allen, J. D., Wrapp, D., McLellan, J. S. & Crispin, M. Site-specific glycan analysis of the SARS-CoV-2 spike. Science eabb9983 (2020) doi:10.1126/science.abb9983.

22. Rothe, C. et al. Transmission of 2019-NCOV infection from an asymptomatic contact in Germany. New England Journal of Medicine vol. 382 970–971 (2020).

23. Liu, C. et al. Viral Architecture of SARS-CoV-2 with Post-Fusion Spike Revealed by Cryo-EM. bioRxiv 1–17 (2020) doi:10.1101/2020.03.02.972927.

24. Chlanda, P. et al. The hemifusion structure induced by influenza virus haemagglutinin is determined by physical properties of the target membranes. Nat. Microbiol. 1, 16050 (2016).

25. Xiong, X. et al. Receptor binding by a ferret-transmissible H5 avian influenza virus. Nature 497, 392–396 (2013).

26. Gui, M. et al. Cryo-electron microscopy structures of the SARS-CoV spike glycoprotein reveal a prerequisite conformational state for receptor binding. Cell Res. 27, 119–129 (2017).

27. Wasilewski, S., Calder, L. J., Grant, T. & Rosenthal, P. B. Distribution of surface glycoproteins on influenza A virus determined by electron cryotomography. Vaccine 30, 7368–7373 (2012).

28. Korber, B. et al. Spike mutation pipeline reveals the emergence of a more transmissible form of SARS-CoV-2. bioRxiv 2020.04.29.069054 (2020) doi:10.1101/2020.04.29.069054.

29. Wan, W. et al. Structure and assembly of the Ebola virus nucleocapsid. Nature 551, 394–397 (2017).

30. Mastronarde, D. N. Automated electron microscope tomography using robust prediction of specimen movements. J. Struct. Biol. 152, 36–51 (2005).

31. Hagen, W. J. H., Wan, W. & Briggs, J. A. G. Implementation of a cryo-electron tomography tilt-scheme optimized for high resolution subtomogram averaging. J. Struct. Biol. 197, 191–198 (2017).

32. Kremer, J. R., Mastronarde, D. N. & McIntosh, J. R. Computer visualization of threedimensional image data using IMOD. J. Struct. Biol. 116, 71–76 (1996).

33. Grant, T. & Grigorieff, N. Measuring the optimal exposure for single particle cryo-EM using a 2.6 Å reconstruction of rotavirus VP6. Elife 4, (2015).

34. Xiong, Q., Morphew, M. K., Schwartz, C. L., Hoenger, A. H. & Mastronarde, D. N. CTF determination and correction for low dose tomographic tilt series. J. Struct. Biol. 168, 378–387 (2009).

35. Turoňová, B., Schur, F. K. M., Wan, W. & Briggs, J. A. G. Efficient 3D-CTF correction for cryo-electron tomography using NovaCTF improves subtomogram averaging resolution to 3.4 Å. J. Struct. Biol. 199, 187–195 (2017).

36. Galaz-Montoya, J. G., Flanagan, J., Schmid, M. F. & Ludtke, S. J. Single particle tomography in EMAN2. J. Struct. Biol. 190, 279–290 (2015).

37. Nickell, S. et al. TOM software toolbox: Acquisition and analysis for electron tomography. J. Struct. Biol. 149, 227–234 (2005).

38. Förster, F., Medalia, O., Zauberman, N., Baumeister, W. & Fass, D. Retrovirus envelope protein complex structure in situ studied by cryo-electron tomography. Proc. Natl. Acad. Sci. U. S. A. 102, 4729–4734 (2005).

39. Kovtun, O. et al. Structure of the membrane-assembled retromer coat determined by cryo-electron tomography. Nature 561, 561–564 (2018).

40. Peukes, J. et al. The native structure of the full-length, assembled influenza A virus matrix protein, M1. bioRxiv 2020.06.24.168567 (2020) doi:10.1101/2020.06.24.168567.

41. Pettersen, E. F. et al. UCSF Chimera – A visualization system for exploratory research and analysis. J. Comput. Chem. 25, 1605–1612 (2004).

42. Qu, K. et al. Structure and architecture of immature and mature murine leukemia virus capsids. Proc. Natl. Acad. Sci. U. S. A. 115, E11751–E11760 (2018).

43. Bharat, T. A. M., Russo, C. J., Löwe, J., Passmore, L. A. & Scheres, S. H. W. Advances in Single-Particle Electron Cryomicroscopy Structure Determination applied to Subtomogram Averaging. Structure 23, 1743–1753 (2015).

44. Scheres, S. H. W. Processing of Structurally Heterogeneous Cryo-EM Data in RELION. Methods Enzymol. 579, 125–57 (2016).

45. Zivanov, J. et al. New tools for automated high-resolution cryo-EM structure determination in RELION-3. Elife 7, (2018).

46. Zheng, S. Q. et al. MotionCor2: anisotropic correction of beam-induced motion for improved cryo-electron microscopy. Nat. Methods 14, 331–332 (2017).

47. Rohou, A. & Grigorieff, N. CTFFIND4: Fast and accurate defocus estimation from electron micrographs. J. Struct. Biol. 192, 216–221 (2015).

48. Bepler, T. et al. Positive-unlabeled convolutional neural networks for particle picking in cryo-electron micrographs. Nat. Methods 16, 1153–1160 (2019).

49. Bai, X., Rajendra, E., Yang, G., Shi, Y. & Scheres, S. H. W. Sampling the conformational space of the catalytic subunit of human γ-secretase. Elife 4, (2015).

50. Emsley, P. & Cowtan, K. Coot: model-building tools for molecular graphics. Acta Crystallogr. Sect. D Biol. Crystallogr. 60, 2126–2132 (2004).

51. Afonine, P. V. et al. Real-space refinement in PHENIX for cryo-EM and crystallography. Acta Crystallogr. Sect. D Struct. Biol. 74, 531–544 (2018).

